# Differential Effects of Ganglioside Lipids on the Conformation and Aggregation of Islet Amyloid Polypeptide

**DOI:** 10.1101/2024.05.26.595964

**Authors:** Samuel D. McCalpin, Lina Mechakra, Magdalena I. Ivanova, Ayyalusamy Ramamoorthy

## Abstract

Despite causing over 1 million deaths annually, Type 2 Diabetes (T2D) currently has no curative treatments. Aggregation of the islet amyloid polypeptide (hIAPP) into amyloid plaques plays an important role in the pathophysiology of T2D and thus presents a target for therapeutic intervention. The mechanism by which hIAPP aggregates contributes to the development of T2D is unclear but are proposed to involve disruption of cellular membranes. However, nearly all research on hIAPP-lipid interactions has focused on anionic phospholipids, which are primarily present in the cytosolic face of plasma membranes. We seek here to characterize the effects of three gangliosides, the dominant anionic lipids in the outer leaflet of the plasma membrane, on the aggregation, structure, and toxicity of hIAPP. Our results show a dual behavior that depends on the molar ratio between the gangliosides and hIAPP. For each ganglioside, a low lipid:peptide ratio enhances hIAPP aggregation and alters the morphology of hIAPP fibrils, while a high ratio eliminates aggregation and stabilizes an α-helix-rich hIAPP conformation. A more negative lipid charge more efficiently promotes aggregation, and a larger lipid headgroup improves inhibition of aggregation. hIAPP also alters the phase transitions of the lipids, favoring spherical micelles over larger tubular micelles. We discuss our results in the context of available lipid surface area for hIAPP binding and speculate on a role for gangliosides in facilitating toxic hIAPP aggregation.

## Introduction

Type 2 Diabetes (T2D) is a top 10 cause of global mortality and accounts for over 1 million deaths annually.^1^ While treatments exist to manage symptoms of T2D, such as insulin resistance and reduced insulin secretion, no cure currently exists.^2^ The prevalence of T2D is on the rise, particularly in developed nations, which emphasizes the pressing need for treatments that target its root cause.^1^ While the pathology of T2D is multifactorial, protein aggregation in the islets of the pancreas plays an important role in reducing insulin-secretion capacity by destroying insulin-secreting β-cells.^3^ T2D belongs to a class of conditions called amyloidosis, in which amyloid plaques composed of fibrillar, β-sheet-rich protein aggregates form in affected tissues.^4^ Islet amyloid primarily comprises aggregates of the islet amyloid polypeptide (IAPP), a 37-residue peptide hormone that is expressed and secreted in response to elevated blood glucose, alongside insulin.^4–8^ A wealth of evidence links IAPP aggregation to the development of T2D. For example, animals that have an aggregating variant of IAPP will spontaneously develop diabetes, while animals with IAPP non-aggregating variants that do not develop diabetes.^4,9–11^ Furthermore, diabetes can be induced in these animals by transgenic modification to express human IAPP (hIAPP).^12,13^ Aggregating variants of IAPP are also toxic to cultured cells while non-aggregating variants are not.^14^ However, the toxicity does not derive from the mature amyloid fibrils.^15–17^ Rather, intermediate aggregates, or “oligomers,” appear to directly cause toxicity to cells.^18^

Consensus has not been reached on a mechanism of IAPP cytotoxicity, but leading theories implicate interactions between IAPP and lipid membranes. IAPP binds to and permeabilizes lipid bilayers containing anionic lipids, so it has been proposed that IAPP oligomers might disrupt plasma membranes.^19,20^ Some studies suggest that IAPP can penetrate cell membranes and form β-barrel pores that act as ion channels, leading to a disruption of membrane integrity, imbalanced cellular homeostasis, and cell death.^21,22^ Alternative hypotheses posit that IAPP oligomers disrupt lipid membranes via a detergent-like mechanism, siphoning lipids from the membrane surface, or by direct interaction with protein receptors in the cell membrane, causing a signaling cascade that results in apoptosis.^23,24^ Regardless, proximity to or direct interaction with cellular membrane lipids appears to mediate IAPP toxicity to cells.

However, most of the research in this area has utilized model membranes containing anionic phospholipids. While phospholipids and cholesterol are the dominant classes of lipids in cell membranes, anionic phospholipids primarily localize in the cytosol-facing membrane leaflets.^25–31^ Thus, these previous studies may not provide sufficient insight into the interaction between IAPP and the outer leaflet of the plasma membrane. This is potentially an issue because intracellular IAPP is stored in secretory granules and thus unlikely to encounter cytosol-facing membrane surfaces.^32^ Instead, IAPP might interact with anionic lipids in the extracellular face of the plasma membrane, such as gangliosides. Gangliosides are glycosphingolipids composed of a ceramide tail and an oligosaccharide headgroup that contains at least one sialic acid moiety.^33^ They are also the major anionic lipid component of the outer leaflet of plasma membranes, comprising 15 mol % of the outer leaflet lipids in red blood cells.^29,34–36^ When cholesterol is abundant in the plasma membrane, gangliosides form clusters known as rafts. Ganglioside-containing lipid rafts have high negative-charge density to attract positively charged membrane-binding proteins, like IAPP.

Gangliosides have been implicated in amyloid formation associated with Alzheimer’s Disease (AD) and Parkinson’s Disease (PD), but relatively little is known about their role in islet amyloidosis.^37,38^ It is notable that there is considerable uncertainty in the ganglioside content of pancreatic cells relevant to T2D. Whole human pancreas extracts mostly contain the gangliosides GM3, GD3, and GD1a.^39^ But human pancreatic islets, which contain the insulin- and IAPP-secreting β-cells, are depleted in GM3, GD3, and GD1a compared to whole pancreas and enriched in an unidentified GM2-comigrating ganglioside.^39^ While rat islet and whole pancreas ganglioside compositions varied greatly between two studies, the islets possessed greater total ganglioside content, relative to whole rat pancreas, suggesting that gangliosides are abundant in pancreatic islets.^40,41^ One study from Wakabayashi et. al. supports the involvement of gangliosides in mediating toxicity associated with aggregated IAPP.^42^ They found that disrupting ganglioside rafts in the cell membrane reduced amyloid formation and toxicity by hIAPP and that hIAPP did not aggregate or cause toxicity when incubated with cells that do not produce gangliosides.^42^ MD simulations similarly observed that monomeric IAPP bound on or near GM3 lipid rafts in a phospholipid membrane and that the GM3 clusters facilitated a helix-to-β-sheet structural conversion of membrane-bound IAPP.^43^

Though there is evidence to implicate gangliosides in islet amyloidosis, to our knowledge there has been no experimental work to define the molecular details of an interaction between gangliosides and hIAPP. As such, we performed a biophysical characterization of hIAPP aggregation, structure, and morphology in the presence of ganglioside-containing membranes. For our studies, we chose to work with three ganglioside lipids – GM1, GM3, and GD3 (**Figure 1**). GM3 and GD3 were selected because they are physiologically relevant to the pancreatic environment and GM1 would allow a comparison to previous research with amyloid-β (Aβ) and α-Synuclein (αS), the amyloid peptides associated with AD and PD, respectively.^37,38,44^ We investigated the effects of each ganglioside on the aggregation behavior of hIAPP with kinetic fluorescence assays, CD spectroscopy, TEM, and cell viability assays. Our results demonstrate a dual, concentration-dependent effect on hIAPP aggregation kinetics, conformation, and toxicity. We then discuss our findings in the context of mechanisms by which gangliosides could mediate IAPP toxicity.

**Figure 1.**
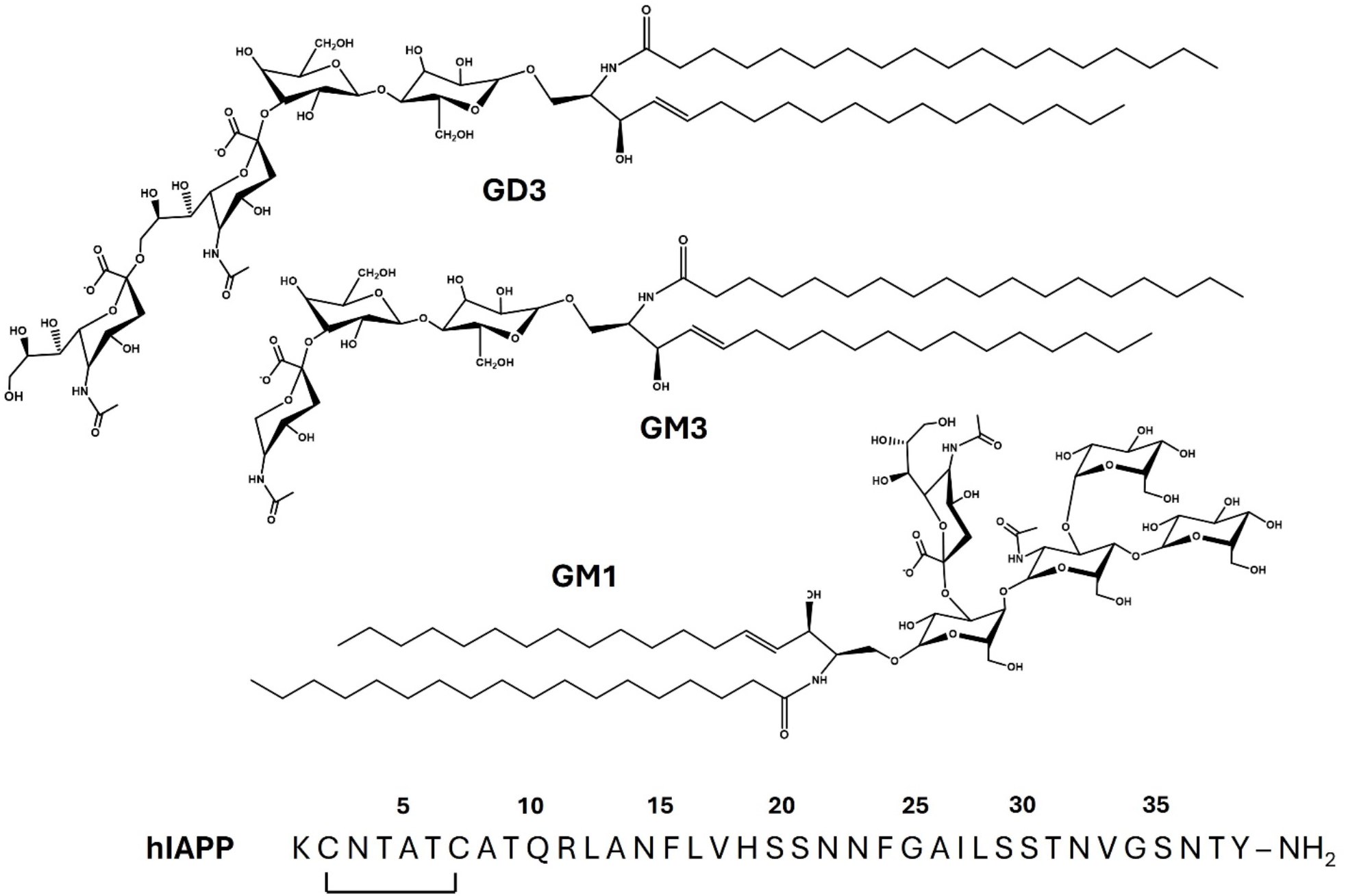
Molecular structures of ganglioside lipids and amino acid sequence of hIAPP. The molecular structures for the noted gangliosides are shown and the amino acid sequence of hIAPP is displayed with the intramolecular disulfide bond and amidated C-terminus noted.

## Methods and Materials

### Materials

C-amidated hIAPP (purity ≥ 95%) was purchased from Anaspec (Fremont, CA, USA). The gangliosides GM1 (ovine brain extract, sodium salt, catalog #860065), GM3 (bovine brain extract, ammonium salt, catalog #860058), and GD3 (bovine brain extract, ammonium salt, catalog #860060) were obtained from Avanti Polar Lipids (Alabaster, AL, USA). All other chemicals were purchased from Sigma Aldrich (St. Louis, MO, USA), except for thioflavin T (ThT), which was purchased from Cayman Chemicals (Ann Arbor, MI, USA).

### Peptide and Lipid Sample Preparation

Prior to use, hIAPP was dissolved in hexafluoro-isopropanol (HFIP) to a concentration of 1 mM and allowed to incubate at room temperature for one hour. hIAPP was aliquoted from this stock and lyophilized. Sample peptide concentrations were calculated based on these aliquots. Lipid stock solutions were prepared at 2.5 mg/mL in a 1:1 mixture of methanol and chloroform. Lyophilized hIAPP and lipid stocks were stored at -20 °C until use. From the stock solutions, lipids were aliquoted, dried to a film under a nitrogen stream, and further dried under a vacuum overnight. These films were resuspended in aqueous buffer for one hour and used immediately.

### Thioflavin T Fluorescence Assay

All samples for the ThT fluorescence assay were prepared on ice, containing a sodium phosphate buffer (10 mM sodium phosphate, 100 mM NaCl, pH 7.4) with 20 µM ThT and the lipid concentrations noted in figures. Lyophilized hIAPP was dissolved in the same phosphate buffer and added to samples immediately prior to beginning each measurement. Samples were added to a black-walled, 384-well plate with a clear, flat bottom (Greiner catalog #07000892) in quadruplicate (30 µL per well). Fluorescence was measured without shaking in a FLUOstar Omega microplate reader (BMG Labtech Inc) by exciting at 440 nm and measuring fluorescence at 490 nm every 8 minutes with gain set at 90%. The temperature inside the microplate reader was maintained at 25 °C for the duration of the experiment. Fluorescence data is shown as the average of the four replicates per sample, with error bars representing one standard deviation. All ThT assays were independently repeated at least once to ensure reproducibility. Amylofit was used to calculate half-times for each trial, and these were averaged and reported with one standard deviation error.^45^

### Transmission Electron Microscopy

Negatively stained specimens for TEM were prepared by applying 5 μL of sample to hydrophilic 400-mesh carbon-coated Formvar support films mounted on copper grids (Ted Pella, Inc., cat# 01702-F). The samples were allowed to adhere for 4 min, washed twice with ddH2O, and stained for 60-90 sec with 5 μL of 1% uranyl acetate (Ted Pella, Inc.). All samples were imaged at an accelerating voltage of 60 kV in a JEM 1400 Plus (JOEL). Images were collected from at least three grid regions at magnifications from 10,000x – 60,000x, and representative micrographs were reported here.

### Circular Dichroism

Circular dichroism (CD) samples were prepared by mixing 50 µM hIAPP with 0, 75, or 750 µM ganglioside lipid in sodium phosphate buffer (10 mM sodium phosphate, 100 mM NaF, pH 7.4). Spectra were measured as an average of 10 accumulations with a Jasco CD spectrophotometer every hour for 24 hours. The sample cell was maintained at 25 °C for the duration of the experiments. Experimental parameters were 100 nm/minute scanning speed, 1 nm bandwidth, 0.5 nm data pitch, 1 s data integration, and 200 mdeg CD scale. Reference spectra were measured using samples without IAPP and automatically subtracted from sample spectra. BestSel was used to estimate secondary structure contents.^46^ For samples with 50 µM hIAPP and 750 µM lipid, secondary structure estimates are reported as the average from all timepoints, with one standard deviation error.

### Cell Toxicity Assay

Toxicity to rat pancreatic β-cells was assessed with the MTT cell viability assay. RIN-5F cells (ATC CRL-2058, batch 61465080) were grown in RPMI-1640 medium with GlutaMAX (diluted from 100x solution Fisher cat#35-050-061), 10% FBS, and Penicillin/Streptomycin at 37 °C and 5% CO_2_. Cells were passaged a minimum of three splitting cycles after revival from frozen stocks prior to toxicity experiments and discarded after 25 passages. The MTT assay was performed with the Promega CellTiter 96 cell proliferation assay kit (Promega G4000). In a transparent 96-well plate, 40,000 cells were added to wells in 90 µL of cell growth medium and allowed to adhere for 24 hours. Samples were then added from 10x stocks, with five replicates per sample, to reach a final hIAPP concentration of 10 µM and final lipid concentrations of 10 µM or 100 µM. The sample plates were incubated for 48 hours at 37 °C and under 5% CO_2_. Per the manufacturer protocol, 15 µL of MTT dye solution was added to each sample, followed by four hours of incubation. Then, 100 µL of stop solution was added to each well. Cell proliferation was assessed by measuring the difference between A_570_ and A_700_ for each well, averaging the replicates for each sample, subtracting the absorbance difference of a sample with 1% SDS, and normalizing relative to a buffer control. This experiment was performed independently three times and cell viability was reported as an average of 15 sample replicates. A one-way ANOVA test was used for statistical analysis.^47^

## Results

### Ganglioside Lipids Exerted a Dual Effect on hIAPP Aggregation

To investigate the effects of ganglioside lipids on hIAPP aggregation, we performed ThT fluorescence assays with 10 µM hIAPP and concentration series of GM1, GM3, and GD3. The ThT fluorescence data revealed two distinct behaviors of gangliosides on hIAPP aggregation (**Figure 2**). At low concentrations, gangliosides reduced the time to half-maximal fluorescence (t_1/2_) and either had no effect or increased the maximum fluorescence intensity (F_max_). For 10 µM hIAPP, the greatest reduction in t_1/2_ was observed with a concentration of 15 µM for GM1 and GD3 and 10 µM for GM3. F_max_ was most increased with 1 µM lipid for GM1 and GM3 and 5 µM lipid for GD3. GD3 decreased t_1/2_ and increased F_max_ to a greater extent than either GM1 or GM3. At higher lipid concentrations, all three gangliosides increased t_1/2_ and reduced F_max_ to the baseline level. Both GM1 and GD3 eliminated the increase in ThT fluorescence with 50 µM lipid, while 100 µM GM3 was required to do the same.

**Figure 2.**
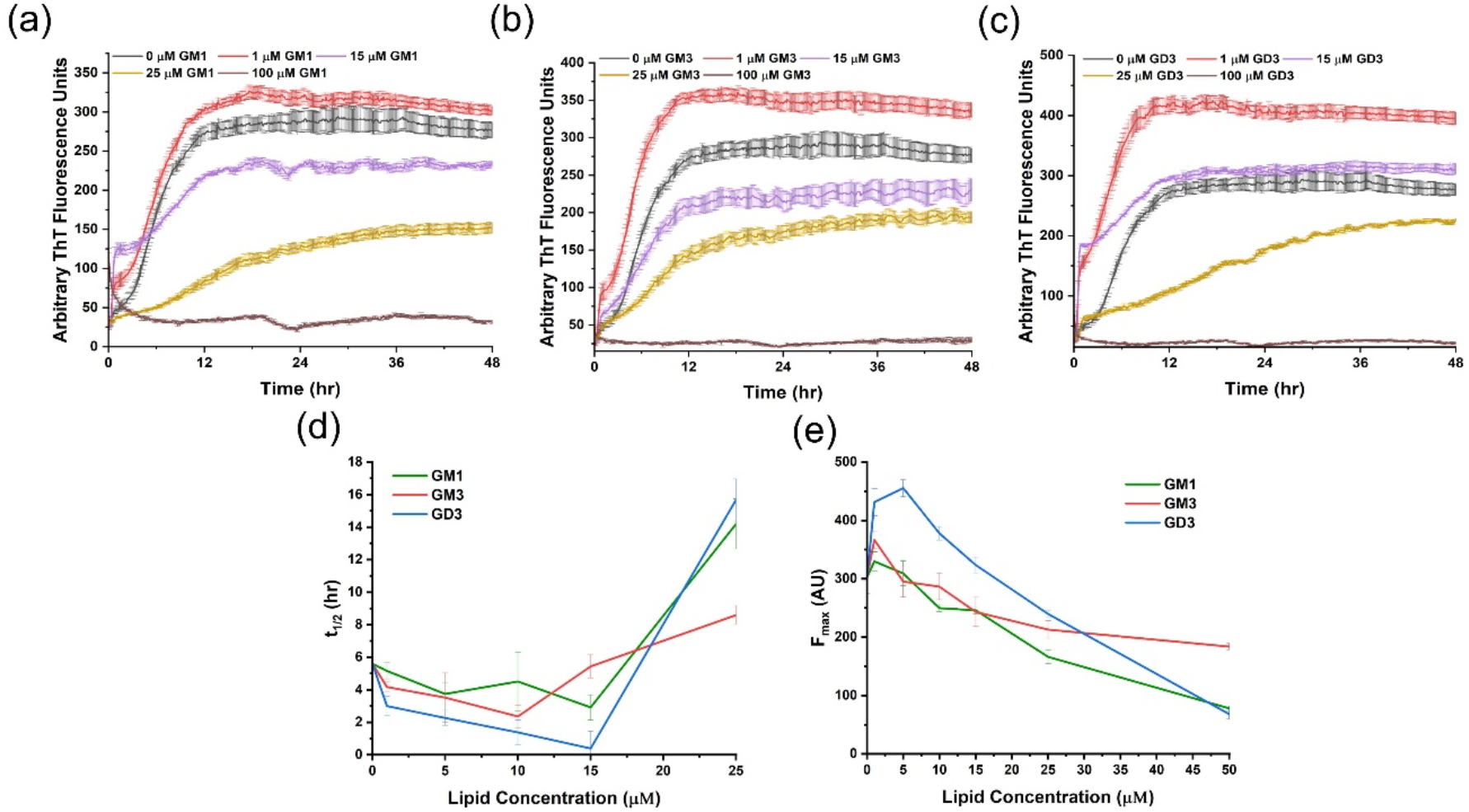
Aggregation kinetics of hIAPP with gangliosides monitored by ThT fluorescence. Samples contained 10 µM hIAPP, 20 µM ThT, 10 mM sodium phosphate, 100 mM NaCl, pH 7.4, and the noted concentrations of **(A)** GM1, **(B)** GM3, and **(C)** GD3. **(D)** Half-times and **(E)** maximum fluorescence intensities were calculated for each condition and plotted versus lipid concentration.

Previous work has measured a GM1 phase transition at a concentration in the range 10-100 µM that has been assigned to the critical micelle concentration (CMC).^48–52^ However, these measurements relied on pyrene fluorescence or triiodide formation assays that report a general increase in hydrophobicity rather than a direct observation of micellar species. Several reports using more direct measurements of particle sizes, such as sedimentation or light scattering, determined a sub-micromolar CMC of GM1.^53–55^We sought to characterize the lipid species present at different concentrations to clarify this discrepancy and the effect of ganglioside phase on hIAPP aggregation. TEM micrographs (**Figure 3**) showed that spherical or discoidal micelles were present with 15 µM of each ganglioside. Increasing the concentrations of GM1 and GM3 to 150 µM resulted in the formation of worm-like or tubular micelles, which potentially explains the lipid phase transition that was measured by pyrene fluorescence and misattributed to the CMC. At this concentration, GM1 formed a mixture of tubular and spherical micelles, while GM3 was entirely tubular. In contrast, the more negatively charged GD3 underwent no phase change between 15-150 µM, remaining entirely as spherical micelles, so it seemed unlikely that the opposite effects of the gangliosides on hIAPP aggregation arose from distinct ganglioside species. Accordingly, further ThT experiments with different concentrations of hIAPP (**Figure 4**) demonstrated that the impact of gangliosides on hIAPP aggregation depended primarily on the lipid:peptide molar ratio rather than the lipid phase. The lipid:peptide molar ratio required to completely inhibit aggregation was greatest for GM3 and least for GM1, but the molar ratio for each lipid varied across replicates, between 4-10x lipid.

**Figure 3.**
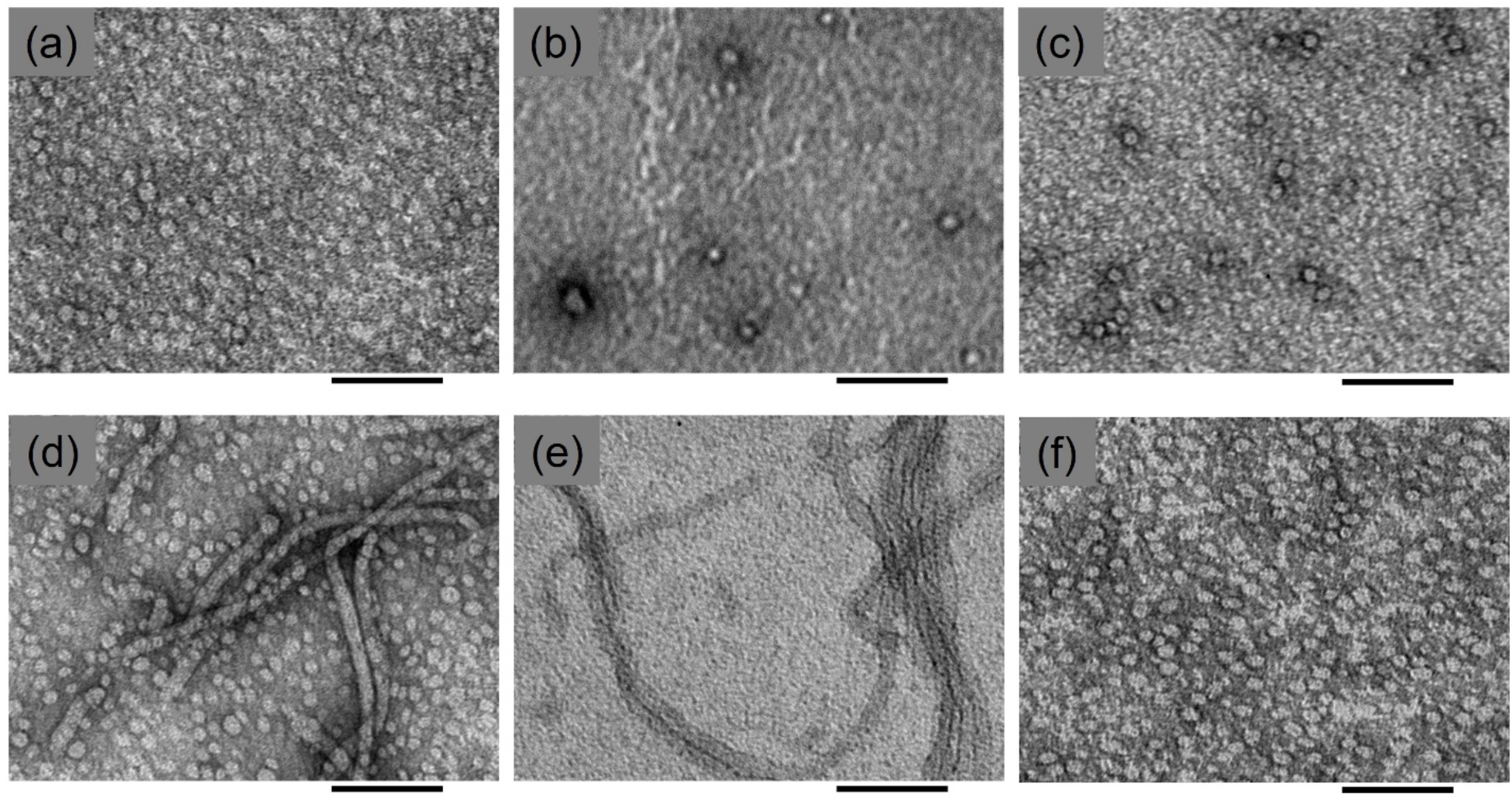
TEM micrographs of gangliosides. Samples were prepared with **(A)** 15 µM GM1, **(B)** 15 µM GM3, **(C)** 15 µM GD3, **(D)** 150 µM GM1, **(E)** 150 µM GM3, or **(F)** 150 µM GD3 in 10 mM sodium phosphate, 100 mM NaCl, pH 7.4. Scale bars = 100 nm.

**Figure 4.**
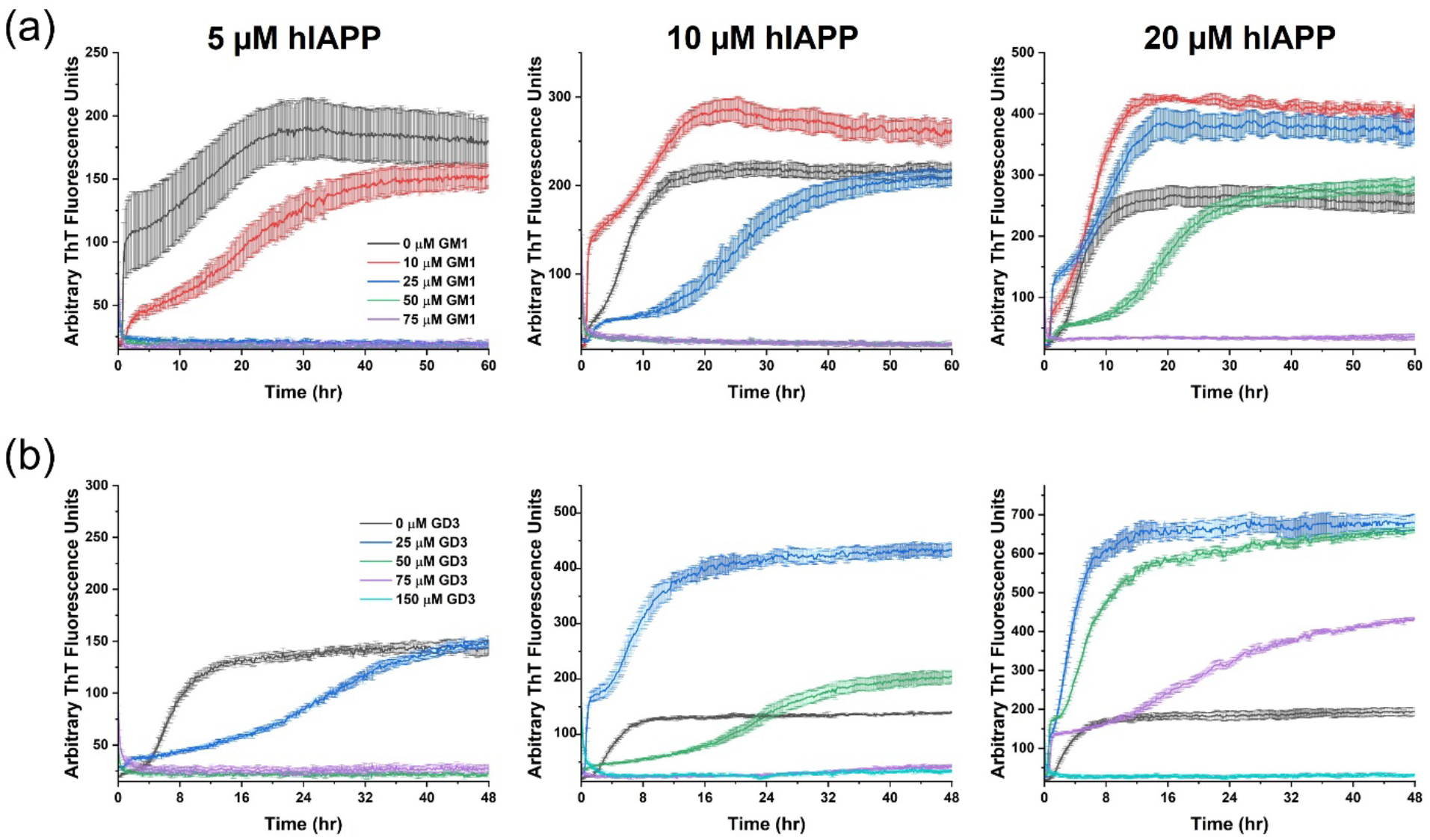
hIAPP aggregation kinetics are dependent on the molar ratio between ganglioside and hIAPP. ThT fluorescence assays were performed with the noted concentrations of hIAPP and **(A)** GM1 or **(B)** GD3 in buffer containing 10 mM sodium phosphate, 100 mM NaCl, pH 7.4. Data for GM3 is in **Figure S1**.

To corroborate the effects of gangliosides on hIAPP aggregation observed by ThT fluorescence, we collected TEM micrographs of hIAPP samples after ThT fluorescence reached plateau (**Figure 5**). hIAPP alone formed homogeneous amyloid fibrils. Addition of 1.5x molar excess ganglioside induced the formation of thinner, straighter fibrils with additional amorphous density wrapped around the fibrils. These wrapped fibrils were morphologically similar with each lipid. Greater heterogeneity was observed amongst the lipid-induced aggregates, with short, ribbon-like assemblies of fibrils and fuzzy amorphous aggregates also present (**Figures 5** and **S2**). Increasing the ganglioside concentrations to a 15x molar excess relative to hIAPP eliminated fibril formation. For each of these samples, only round micelles were observed, despite GM1 and GM3 alone forming tubules at the same concentration.

**Figure 5.**
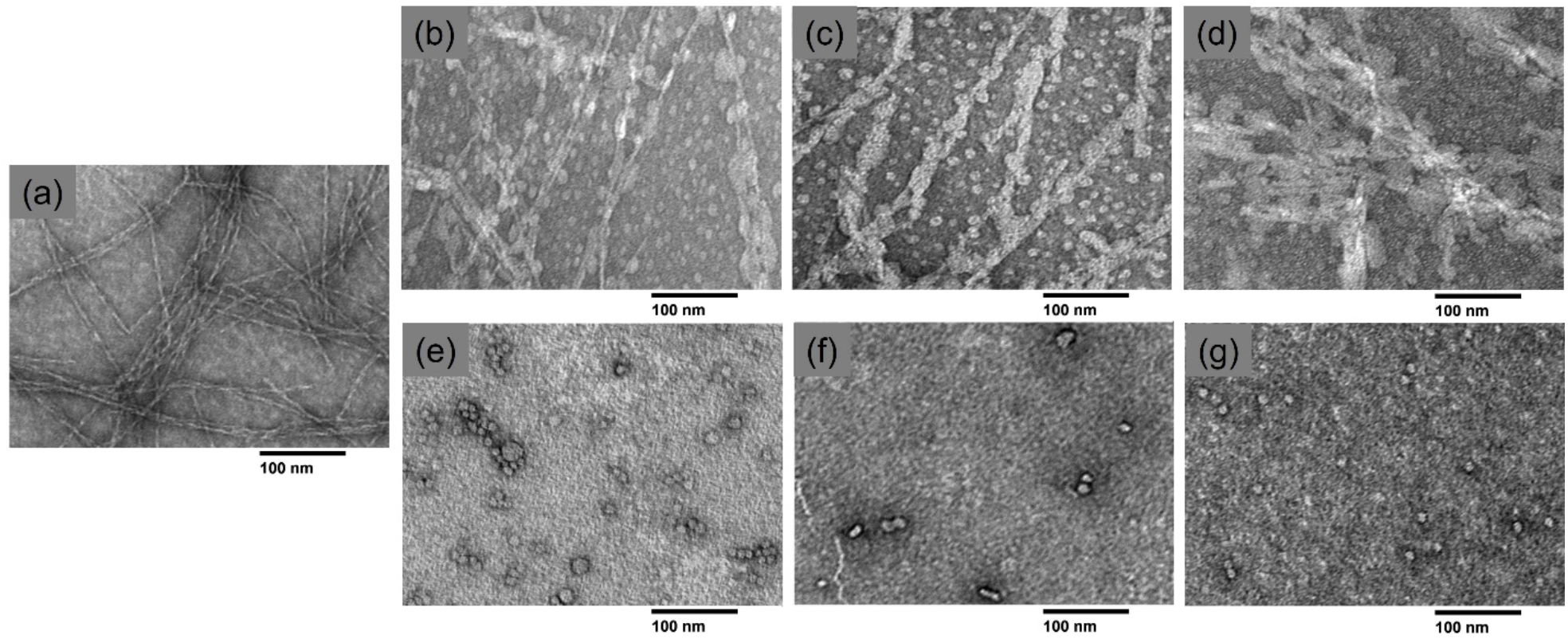
TEM micrographs of hIAPP incubated with gangliosides. Samples containing 10 µM hIAPP and **(A)** no lipids, **(B)** 15 µM GM1, **(C)** 15 µM GM3, **(D)** 15 µM GD3, **(E)** 150 µM GM1, **(F)** 150 µM GM3, **or (G)** 150 µM GD3 were monitored by ThT fluorescence for 96 hr to confirm complete aggregation and stained with 1% uranyl acetate for TEM imaging.

### Gangliosides Induced Conformational Changes in hIAPP

To investigate conformational changes of hIAPP in the presence of gangliosides, we followed the time course of hIAPP aggregation in fibril-seeding (1.5x molar excess ganglioside) and fibril-inhibiting (15x molar excess ganglioside) conditions with CD spectroscopy (**Figure 6**). The CD spectrum of hIAPP alone initially displayed a negative minimum at 201 nm which shifted to 219 nm within a few hours, consistent with a transition from random coil to β-sheet. In contrast, the addition of a 1.5x molar excess of any of the gangliosides resulted in double minima at 208 nm and 221 nm, indicative of α-helix. Over time, these two minima transformed to a single minimum at 219 nm, indicating an α-helix to β-sheet structural conversion. A higher concentration of gangliosides (15x molar excess) produced a similar starting CD spectrum that was stable for at least 24 hours (**Figures 6E** and **S3**). We then deconvoluted the CD spectra of hIAPP with 15x lipid to estimate the secondary structure content of stable lipid-bound hIAPP (**Tables S1** and **S2**). The estimates indicated that, regardless of the ganglioside, the micelle-bound hIAPP mostly contained α-helix and random coil secondary structures. The CD deconvolutions also predicted smaller amounts of β-sheet and turn structures that depended on the ganglioside identity. However, β-sheet and turn content varied substantially based on the range of the CD data used in secondary structure estimations.

**Figure 6.**
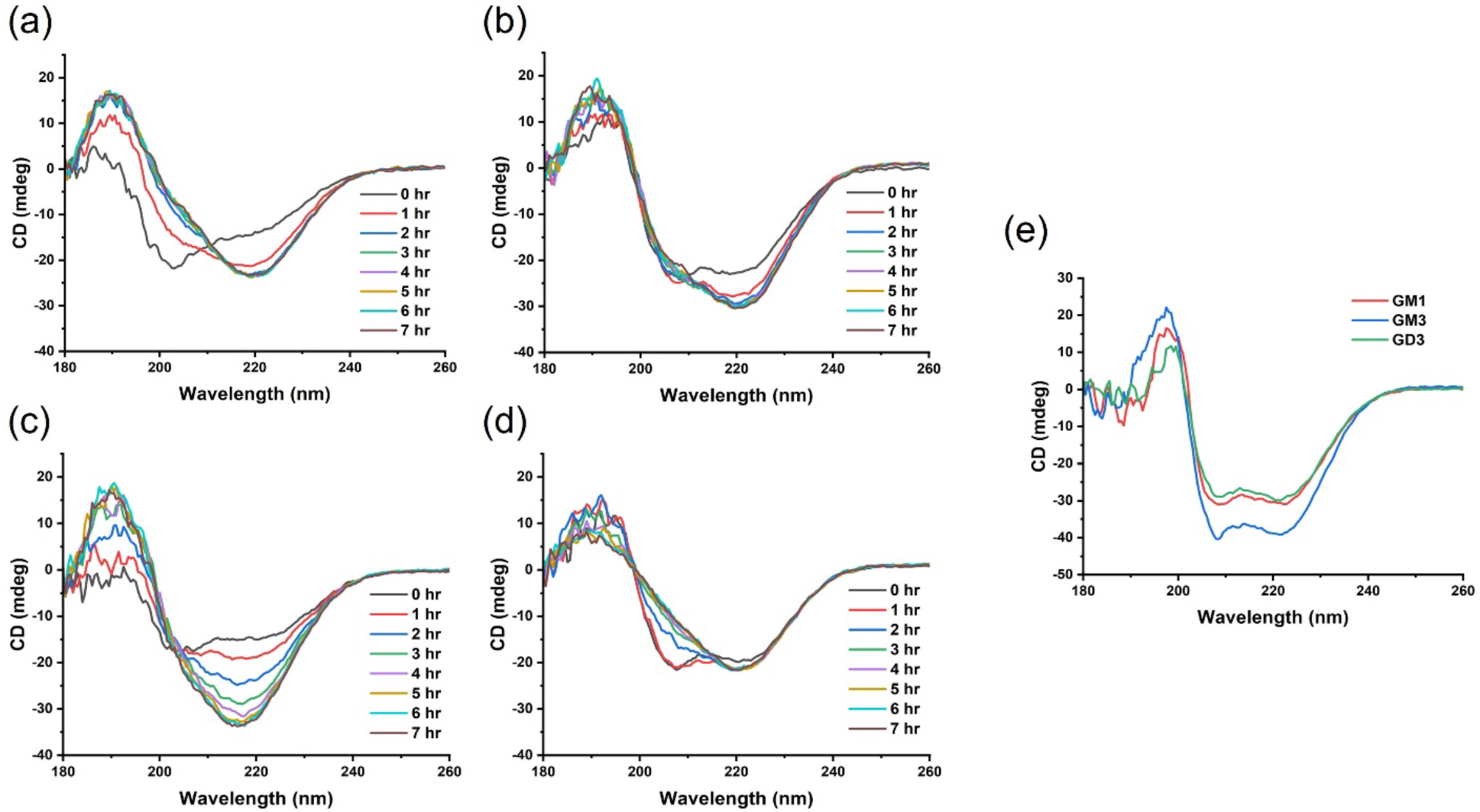
Time-dependent CD spectra of hIAPP incubated with gangliosides. hIAPP (50 µM) was incubated with **(A)** no lipids, **(B)** 75 µM GM1, **(C)** 75 µM GM3, **(D)** 75 µM GD3, or **(E)** 750 µM of each ganglioside in a sodium phosphate buffer (10 mM sodium phosphate, 100 mM NaF, pH 7.4). Spectra were measured every hour for at least 20 hours and are presented for **(A-D)** the first eight hours or **(E)** the first hour.

### Cytotoxicity of Ganglioside-Induced hIAPP Aggregates

We incubated hIAPP and ganglioside lipids with RIN-5F rat insulinoma cells and performed MTT cell viability assays to investigate the connection between ganglioside-induced conformational changes of hIAPP and cytotoxicity (**Figure 7**). hIAPP alone was toxic after 48 hr incubation, reducing cell viability by 57% compared to the buffer control. All the ganglioside concentrations in the absence of hIAPP also decreased cell viability, except for 10 µM GM1 which caused a statistically insignificant reduction of cell viability. At a concentration of 100 µM, GD3 and GM1 were more toxic on their own than GM3, and the ganglioside toxicity was dose-dependent, increasing with concentration. Compared to the ganglioside alone conditions, addition of hIAPP further reduced cell viability for treatments with equimolar (10 µM) of any ganglioside and 10x (100 µM) GM3 but not for treatments with 10x (100 µM) GM1 or GD3. However, we cannot definitively conclude that the high ganglioside concentrations reduced the toxicity of hIAPP because the lipids were toxic themselves, and we could not distinguish between toxicity from hIAPP and toxicity from the gangliosides, particularly for GM1 and GD3.

**Figure 7.**
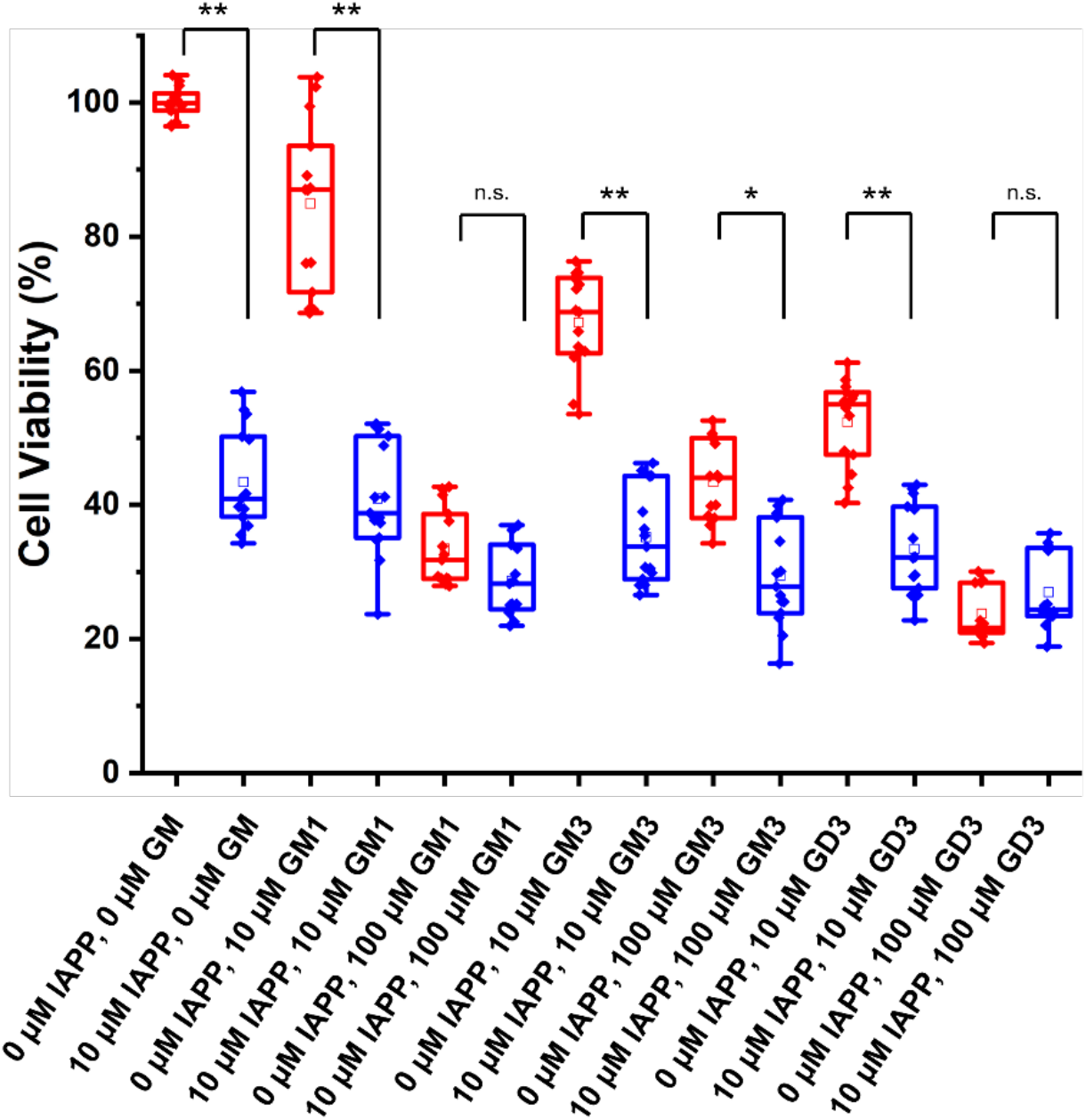
Viability of rat insulinoma cells incubated with hIAPP and gangliosides. Cell viability was measured by the MTT cell viability assay for RIN-5F cells incubated with the noted samples for 48 hr. The box and whisker plots show the sample median (solid line), mean (square), and scatter plot of raw data. A one-way ANOVA test was used for statistical analyses. Pairwise differences between samples with (blue) and without (red) 10 µM hIAPP are denoted as p>0.05 (n.s.), 0.01< p<0.05 (*), or p<0.01 (**). All treatments except for 0 µM hIAPP, 10 µM GM1 (box 3) caused significantly reduced viability compared to the buffer control (box 1).

## Discussion

Biophysical studies have extensively characterized hIAPP interactions with model lipid membranes containing various phospholipids.^19,56–62^ hIAPP binds bilayers composed of DOPG or POPG but not those consisting of DOPC or POPC, indicating a strong preference for negative charge density.^56,57^ Membranes composed of POPS or DOPS also strongly bind hIAPP, confirming that this binding depends primarily on charge and not headgroup identity.^61–63^ Upon binding to phospholipid membranes or micelles, hIAPP adopts a partially α-helical structure.^57,61–68^ A low concentration of anionic phospholipids promotes hIAPP fibrillation while a high concentration is inhibitory.^56,62^ Similarly, vesicles with both zwitterionic and anionic lipids promote fibrillation with low anionic lipid content and inhibit with high anionic lipid content, suggesting that the effect of anionic lipids on hIAPP aggregation is independent of the presence of zwitterionic lipids.^56,62^ The amount of anionic lipid required to inhibit hIAPP aggregation increases with ionic strength, consistent with an electrostatic lipid-peptide interaction.^62^

We determined that the ganglioside lipids GM1, GM3, and GD3 exerted similar effects on hIAPP aggregation, dependent on the lipid:peptide ratio. As with the anionic phospholipids, the gangliosides promoted aggregation with low lipid:peptide but inhibited aggregation with high lipid:peptide. GD3, the most negatively charged ganglioside, most effectively promoted hIAPP aggregation, consistent with an electrostatic model of interaction. Similarly, the gangliosides induced α-helix structure in hIAPP that converted to β-sheet over time with low lipid:peptide and was stable with high lipid:peptide. These opposing effects could be explained by considering the available lipid surface area for binding by hIAPP (**Figure 8**). At high lipid concentrations, more liposomes or micelles provide greater available lipid surface area for hIAPP binding. As a result, hIAPP can bind more diffusely and is thus less likely to self-associate. Reducing the lipid concentration reduces the available binding area for hIAPP, thereby increasing spatial proximity of lipid-bound hIAPP monomers and facilitating their interaction. GM3 required the highest lipid:peptide ratio to inhibit hIAPP aggregation, GD3 the next highest, and GM1 the least. This is in order of the number of headgroup carbohydrate moieties, supporting the idea that lipid surface area facilitates their inhibition of hIAPP aggregation.

**Figure 8.**
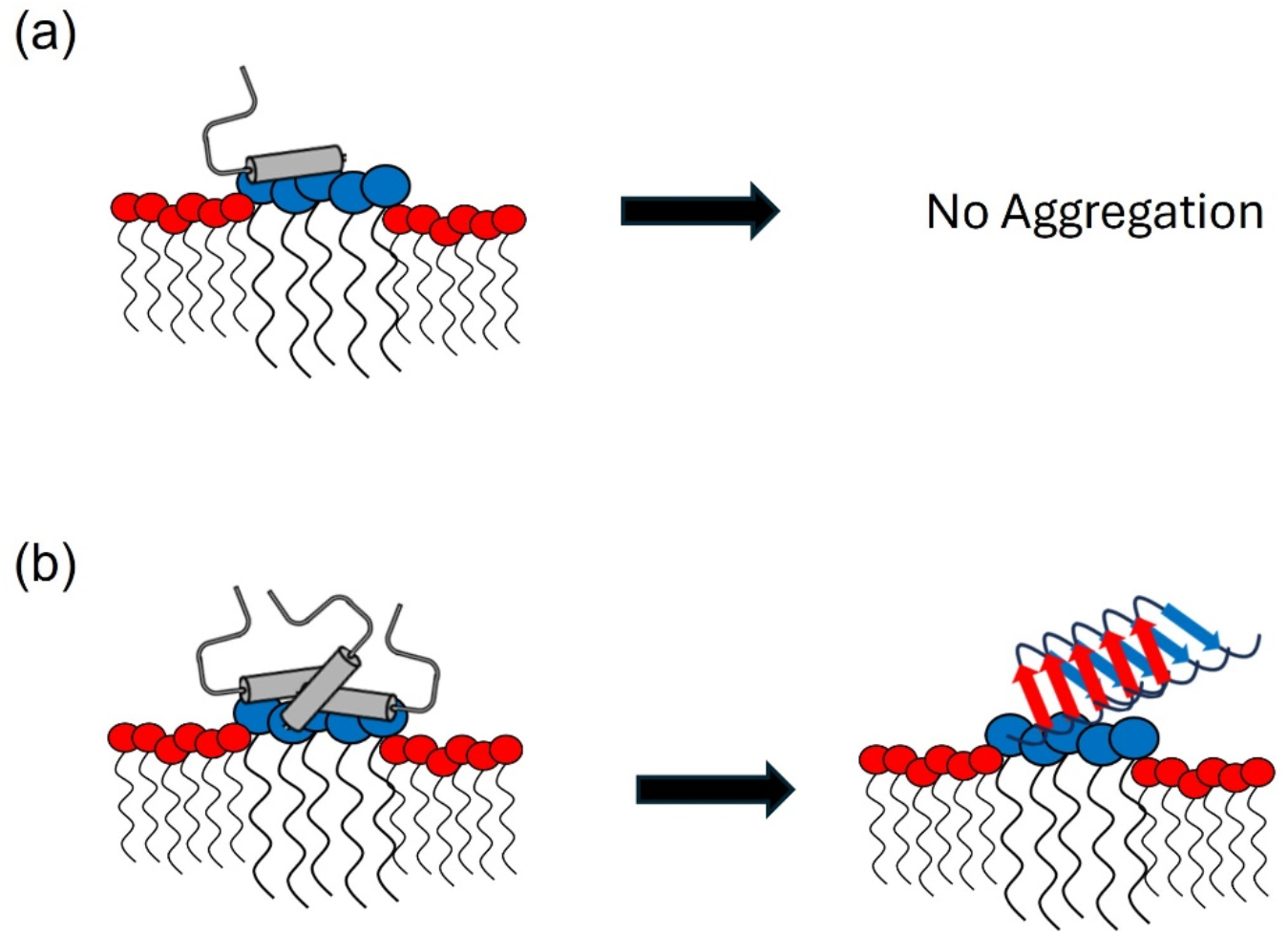
Schematic model for concentration-dependent effects of gangliosides on hIAPP aggregation. **(A)** With a high ratio of ganglioside:peptide, there is sufficient area of ganglioside rafts (blue) in phospholipid (red) membranes for hIAPP monomers (grey) to bind diffusely, preventing aggregation. **(B)** In contrast, a low ganglioside:peptide ratio forces hIAPP monomers to bind in proximity to other monomer on scarce ganglioside rafts, facilitating self-association for amyloid fibril formation.

Using TEM, we also observed that the gangliosides altered the morphology of hIAPP fibrils. In the presence of each ganglioside, an amorphous density formed along the hIAPP fibril length. Fibrils with a similar morphology have been observed following hIAPP aggregation on phospholipid membranes, and the amorphous density has been attributed to lipids coating the fibril surface, though this lipid wrapping has not always been observed under such conditions.^56,69,70^ hIAPP also modified the aggregation of the lipids, preventing formation of worm-like tubules of GM1 and GM3 at high lipid concentrations. Previous work has shown that hIAPP senses membrane curvature and remodels membranes.^71,72^ For instance, hIAPP transformed large POPS vesicles into smaller liposomes, like what we reported.^71^ These observations may support mechanisms by which hIAPP can disrupt ganglioside-containing plasma membranes, leading to cell death. Lipid-wrapped fibrils suggest that hIAPP aggregates could sequester lipids from the membrane, as in the proposed detergent-like mechanism of membrane disruption by hIAPP.^21^ Alternatively, hIAPP might generate local regions of high curvature in the cell membrane, causing stress that ultimately results in perforation of the cell.^72^

In summary, we studied the effects of the gangliosides GM1, GM3, and GD3 on hIAPP aggregation. For each ganglioside, equimolar or lower concentrations relative to hIAPP promoted aggregation, and GD3 was more effective than GM1 or GM3. On the other hand, higher ganglioside concentrations, relative to hIAPP, inhibited hIAPP aggregation. GM1 and GD3 were more efficient inhibitors than GM3, likely due to larger headgroups providing greater surface area for hIAPP binding. We also described changes in hIAPP and ganglioside aggregate morphologies that potentially support curvature-strain-induced and detergent-like mechanisms of hIAPP-induced membrane penetration. However, more work is needed to clarify the molecular underpinnings of membrane disruption by hIAPP. Thus, further research should expand the scope of this report to include physiologically relevant model membranes with zwitterionic phospholipids, cholesterol, and ganglioside rafts. More work is also needed to elucidate the nature and structure of hIAPP aggregates that form in the presence of ganglioside-enriched membranes and how these species correlate with cytotoxicity associated with T2D. Such studies could provide a fuller picture of the toxic nature of hIAPP aggregates and inspire development of more effective therapeutics against T2D.

## Acknowledgements

Funding was provided by National Institutes of Health Grant DK13221401 to A.R.

