## Supporting Information for "Differential Effects of Ganglioside Lipids on the Conformation and Aggregation of Islet Amyloid Polypeptide"

AUTHOR INFORMATION

**Corresponding Authors**

*

*

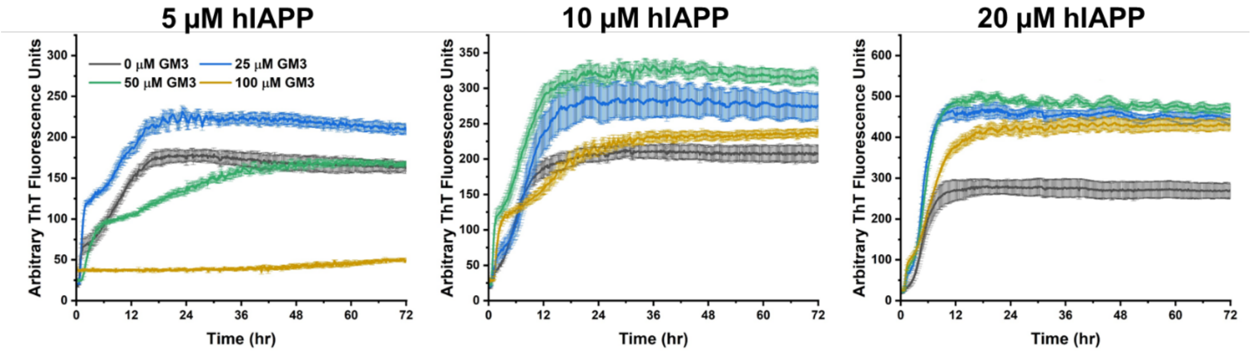


**Figure S1.** hIAPP aggregation kinetics are dependent on the molar ratio between GM3 and hIAPP. ThT fluorescence assays were performed with the noted concentrations of hIAPP and GM3.

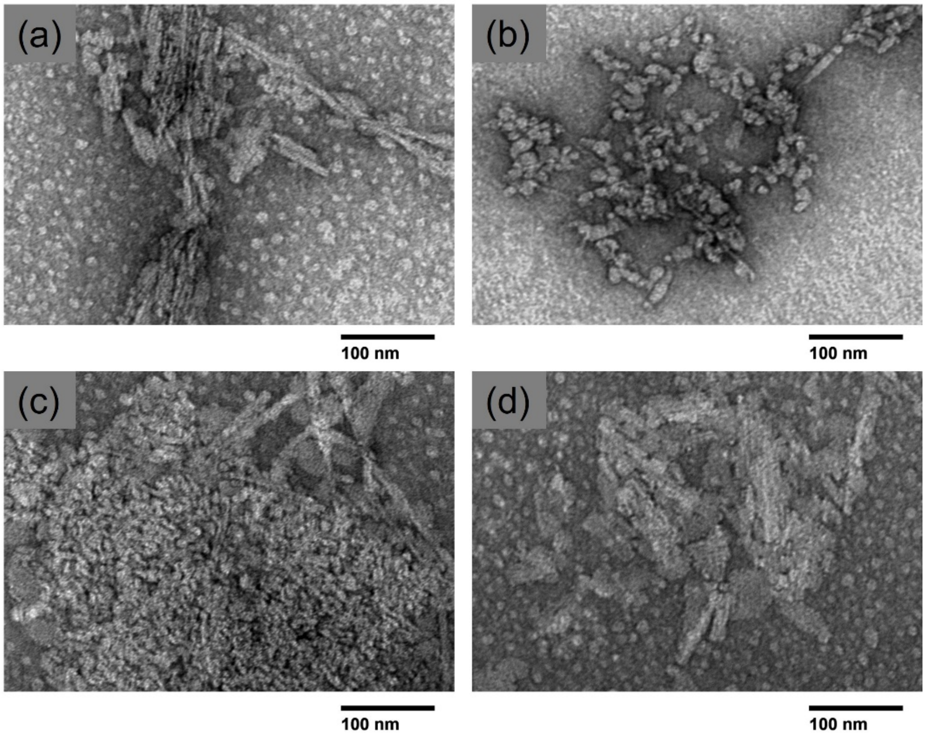


**Figure S2.** TEM micrographs of polymorphic hIAPP-ganglioside species. Samples containing 10 μM hIAPP and **(A)** 10 μM GM1, **(B)** 10 μM GM3, or **(C,D)** 10 μM GD3 were monitored by ThT fluorescence for 96 hr to confirm complete aggregation and negatively stained with 1% uranyl acetate on a copper coated grid for analysis by TEM. Bars representing 100 nm are included beneath each image.

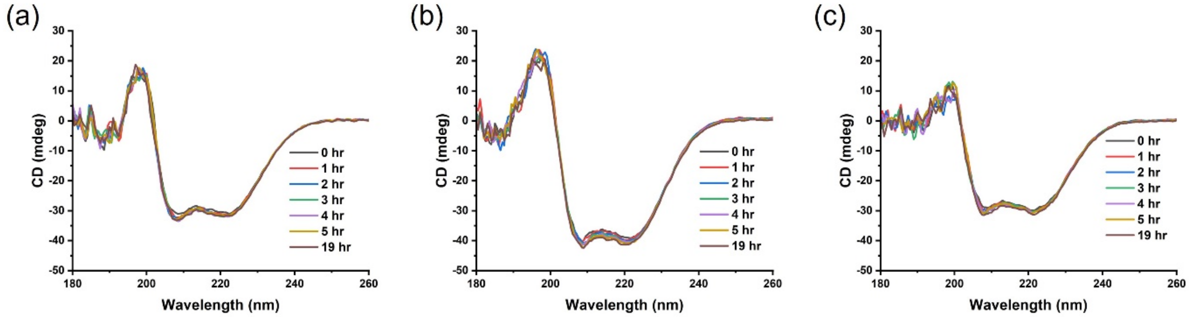


**Figure S3.** CD spectra of hIAPP with 15x gangliosides are stable over time. hIAPP (50 µM) was incubated with 750 µM **(A)** GM1, **(B)** GM3, or **(C)** GD3.

**Table S1.** Secondary structure estimates of micelle-bound hIAPP using CD data from 180-250 nm. Ganglioside samples contained 750 µM lipid, and secondary structure was predicted from previously reported structures of hIAPP bound to SDS micelles (PDB 2KB8) and rat IAPP bound to DPC micelles (PDB 2KJ7). Error is reported as one standard deviation.

| Component % | IAPP + GM1 | IAPP + GM3 | IAPP + GD3 | 2KB8 | 2KJ7 |
|---|---|---|---|---|---|
| **α-helix** | 44.5 ± 0.7 | 50.4 ± 0.7 | 40.5 ± 1.0 | 56.8 | 45.9 |
| **β-sheet** | 10.5 ± 0.7 | 3.1 ± 0.9 | 11.7 ± 0.8 | 0.0 | 0.0 |
| **Turn** | 5.6 ± 0.3 | 0.2 ± 0.3 | 8.2 ± 0.6 | 24.3 | 27.0 |
| **Random Coil** | 39.4 ± 0.8 | 46.2 ± 1.3 | 39.6 ± 0.8 | 18.9 | 27.0 |

**Table S2.** Secondary structure estimates of micelle-bound hIAPP using CD data from 200-250 nm. Ganglioside samples contained 750 μM lipid, and secondary structure was predicted from previously reported structures of hIAPP bound to SDS micelles (PDB 2KB8) and rat IAPP bound to DPC micelles (PDB 2KJ7). Error is reported as one standard deviation.

| Component % | IAPP + GM1 | IAPP + GM3 | IAPP + GD3 | 2KB8 | 2KJ7 |
|---|---|---|---|---|---|
| **α-helix** | 60.5 ± 6.5 | 72.8 ± 6.4 | 50.3 ± 5.6 | 56.8 | 45.9 |
| **β-sheet** | 0.7 ± 1.5 | 1.1 ± 1.8 | 7.1 ± 3.8 | 0.0 | 0.0 |
| **Turn** | 16.0 ± 2.9 | 13.9 ± 1.3 | 16.4 ± 2.1 | 24.3 | 27.0 |
| **Random Coil** | 22.8 ± 8.6 | 12.2 ± 8.9 | 26.2 ± 9.8 | 18.9 | 27.0 |
